# Silica-mediated exacerbation of inflammatory arthritis: A novel murine model

**DOI:** 10.1101/2025.01.07.631488

**Authors:** Lisa M.F. Janssen, Caroline de Ocampo, Dwight H. Kono, Steven Ronsmans, Manosij Ghosh, Peter H.M. Hoet, K. Michael Pollard, Jessica M. Mayeux

**Author notes:** **Correspondence** K. M. Pollard, PhD, Department of Immunology and Microbiology, MB-120, The Scripps Research Institute, 10550 North Torrey Pines Road, La Jolla, CA 92037, USA., J. M. Mayeux, PhD, Department of Immunology and Microbiology, MB-120, The Scripps Research Institute, 10550 North Torrey Pines Road, La Jolla, CA 92037, USA.

## Abstract

**Objective:** The mucosal origin hypothesis in rheumatoid arthritis (RA) posits that inhalant exposures, such as cigarette smoke and crystalline silica (c-silica), trigger immune responses contributing to disease onset. Despite the established risk posed by these exposures, the mechanistic link between inhalants, lung inflammation, and inflammatory arthritis remains poorly understood, partly from the lack of a suitable experimental model. As c-silica accelerates autoimmune phenotypes in lupus models and is a recognized risk factor for several autoimmune diseases, we investigated whether c-silica exposure could induce RA-like inflammatory arthritis in mice.

**Methods:** Two arthritis-prone mouse strains, BXD2/TyJ and HLA-DR4 transgenic (DR4-Tg), were exposed to c-silica or PBS via oropharyngeal instillation. Arthritis was evaluated by clinical signs and histopathology. Autoimmunity was further evaluated by serological analysis, including autoantibodies and cytokines and chemokines. Lung pathology was evaluated by histopathology and immunofluorescent staining for lymphocyte and macrophages.

**Results:** C-silica exposure induced chronic pulmonary silicosis in all mice. In BXD2 mice, this was associated with rapid arthritis development, marked by synovitis, bone erosion, and elevated serum autoantibody levels targeting various antigens, including snRNP and citrullinated protein. Additionally, BXD2 mice exhibited inducible bronchus-associated lymphoid tissue (iBALT) formation and elevated autoantibodies in bronchoalveolar lavage fluid (BALF). Conversely, DR4-Tg mice had no significant arthritis, negligible autoantibody responses, and milder lung inflammation lacking iBALT.

**Conclusion:** We introduce a novel model of c-silica-mediated inflammatory arthritis, creating a novel platform to unravel the molecular and cellular underpinnings of RA and advance understanding of the mucosal origin hypothesis.

**Graphical abstract:** 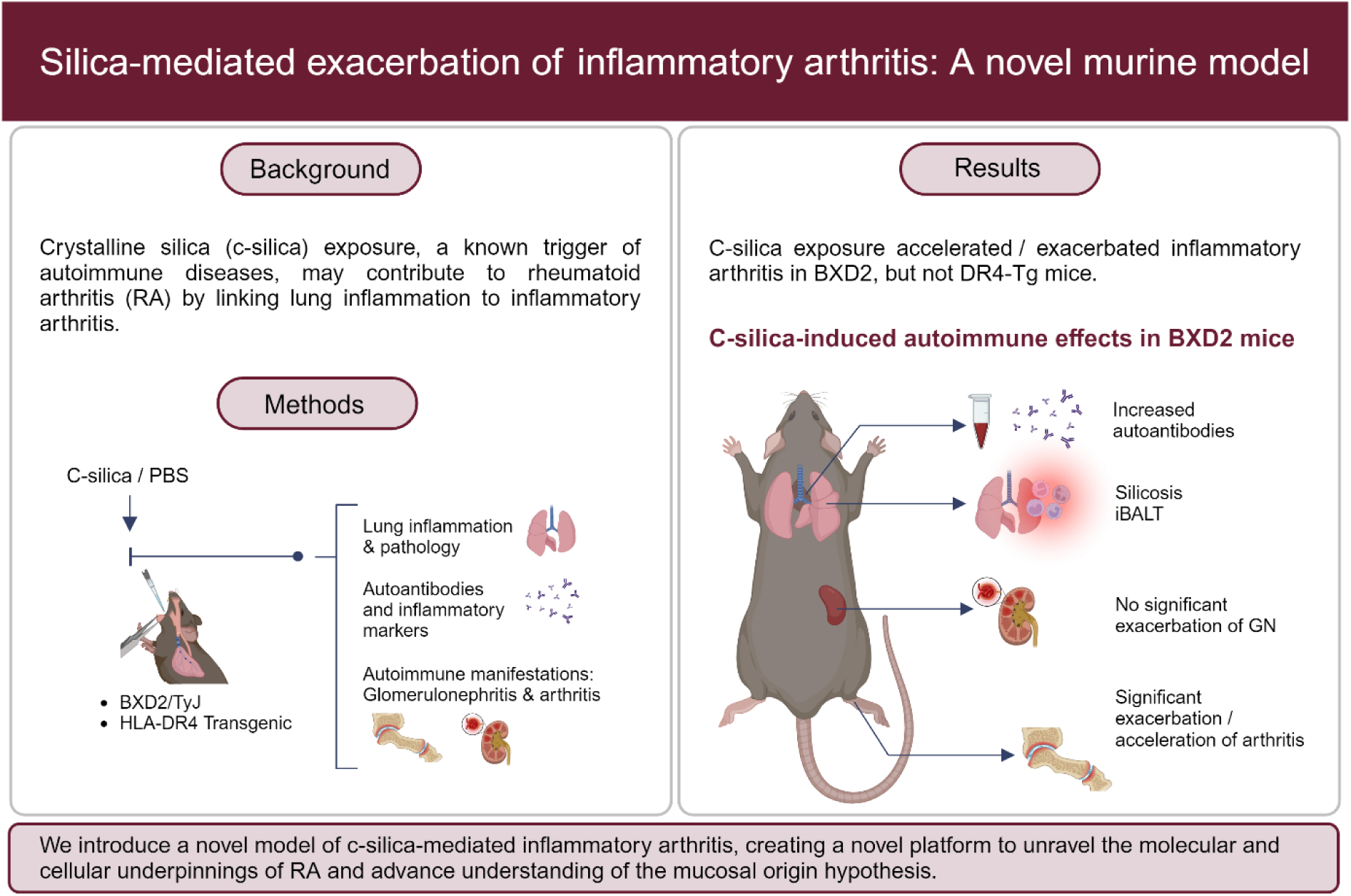

## Introduction

Rheumatoid arthritis (RA) is a chronic autoimmune disease driven by complex interactions between genetic predisposition and environmental exposures, culminating in chronic inflammatory synovitis and joint destruction [1–3]. RA affects ∼0.5 - 1% of the general population, with women being two to three times more susceptible [4]. Among the genetic factors, the HLA-DRB1 “shared epitope” (SE) alleles [5], including HLA-DRB1*0401 (DR4) [6], represent the most significant risk, contributing an estimated 30% of the total genetic component [7].

Environmental risk factors also play a pivotal role in RA pathogenesis, with cigarette smoke and crystalline silica (c-silica), the most well-established triggers for anti-citrullinated protein antibody (ACPA)-positive RA [8, 9]. C-silica exposure alone is associated with a greater than two-fold increase in RA risk [10]. This ubiquitous mineral, found in the Earth’s crust, becomes airborne during occupational activities such as sandblasting, mining, or artificial stone fabrication [11, 12]. Inhalation of respirable c-silica particles causes silicosis, a debilitating nodular inflammatory fibrotic pneumoconiosis [12]. Beyond its impact on pulmonary health, c-silica exposure is associated with several systemic autoimmune diseases in addition to RA, including systemic lupus erythematosus (SLE) and systemic sclerosis (SSc) [13–15].

Using animal models, we [16, 17] and others [18, 19] (reviewed in [20]) have shown that c-silica can provoke autoantibody production and features of systemic autoimmune diseases, such as glomerulonephritis (GN), in certain mouse strains. However, despite its established association with RA [21], no animal models have demonstrated a direct causal link between c-silica exposure and the hallmark inflammatory arthritis of RA. This gap has limited our understanding of the molecular and cellular mechanisms linking inhalant exposures to RA pathogenesis.

Existing RA models have successfully replicated inflammatory and erosive arthritis and are valuable for studying disease progression and evaluating therapeutics [22]. However, these models often rely on induction methods unrelated to known environmental triggers of human disease. To address this limitation, we examined the effects of c-silica exposure in two arthritis-susceptible strains, the DR4-IE transgenic (Tg) (DR4-Tg) and the BXD2/TyJ mice. DR4-Tg mice, which express the RA-associated HLA-SE, develop arthritis when immunized with citrullinated fibrinogen [23], reflecting the relationship between the HLA-SE, ACPA, and arthritis [24, 25]. Since silica nanoparticles elicit protein citrullination [26], we hypothesized that c-silica exposure would lead to ACPA production and subsequent arthritis. Contrastingly, BXD2/TyJ mice develop generalized systemic autoimmunity characterized by antinuclear antibodies (ANA), rheumatoid factor (RF), glomerulonephritis (GN), and erosive inflammatory arthritis [27]. As c-silica exposure aggravates disease in mice susceptible to systemic autoimmunity [19], we further hypothesized it would accelerate and intensify arthritis in this strain. Thus, this study seeks to bridge a critical knowledge gap by leveraging c-silica exposure in genetically susceptible mouse models to elucidate the environmental contributions to RA pathogenesis, focusing on the mucosal origin hypothesis.

## Materials and methods

### Mice

BXD2/TyJ mice (BXD2/TyJ; Jackson Labs Strain #000075) were obtained from The Jackson Laboratory (Bar Harbor, ME) (hereafter referred to as BXD2). HLA-DRB1*0401 (DR4)-transgenic mice (HLA-DR4-IE; Taconic Model #4149) on C57BL/6 background were obtained from Taconic Biosciences (Germantown, NY) (hereafter referred to as DR4-Tg). Number of animals used for each conditions can be found in Supplementary File 1. Mice were housed under specific pathogen-free conditions, with rooms at 20-26°C, 30-70% humidity, and a 12h/12h light-dark cycle. Sanitized cages were replaced bi-weekly with fresh water and food (Teklad 2018, Envigo, Madison, WI, USA) ad libitum. Experiments were initiated in 8-12-week-old male and female animals. All procedures were approved by the Institutional Animal Care and Use Committee at Scripps Research (protocol #08-0150), and use of c-silica was approved by TSRI Department of Environmental Health and Safety.

### Exposure to crystalline silica

C-silica (SiO2 , Min-U-Sil-5, average particle size 1.5-2μm; U.S. C-silica Company, Frederick, MD) was acid washed (1 M HCl, 100°C, 2h), washed (3x) with sterile water, autoclaved (1h, 121°C), dried, and reconstituted in PBS [28]. Immediately prior to use, c-silica was disbursed by sonication, using a sonicator bath. Mice were exposed to a single dose of 5 mg by transoral/oropharyngeal instillation in a volume of 50 µl in PBS, as previously described [17]. Control animals were exposed to 50 µl of PBS. The selected dose of 5 mg in 50 µl provides the most consistent response in the lungs [17, 29]. A single instillation was used based on research indicating a correlation between the intensity of c-silica exposure and the onset of autoimmune responses [15].

### Harvest and tissue collection

Blood for serum was collected from the retro-orbital sinus pre-c-silica exposure and at intermediate time points post-exposure (4, 8, 12 and 20 weeks post-exposure). BXD2 mice were euthanized 1, 2, 6, 12 or 20-weeks post-exposure. DR4-Tg mice were euthanized 20 or 40 weeks post-exposure. Time points were chosen based on autoantibody responses in intermediate timepoint serum samples and visual assessment of arthritis development. During harvest, bronchoalveolar lavage fluid (BALF) was collected and blood for serum was collected from the heart. Spleen and tracheobronchial lymph node (TBLN) weights were determined. Spleen, kidney, TBLN, and lungs were fixed for 48 hours in zinc formalin and transferred to 70% isopropanol. Ankles were fixed for 72 hours in zinc formalin and decalcified in formic acid. Paraffin sections (4 μm) were prepared and stained by the Microscopy and Histology Core Facilities of the La Jolla Institute for Immunology (LJI, La Jolla, CA).

### C-silica-induced lung inflammation and silicosis

Hematoxylin and eosin (H&E) paraffin sections (4 μm) of lungs were prepared by the Microscopy and Histology Core Facilities (LJI, La Jolla, CA), and slides scanned by Leica Aperio AT2 whole slide scanner. Silicosis was scored under blinded conditions for the percent of each lobe affected by alveolitis or peribronchial and perivascular inflammatory infiltrates (5 lobes, 500 maximum score each), as described previously [17]. The values for alveolitis and for peribronchitis- and vasculitis were combined to give a total lung score (max 1,000).

### Fluorescent immunohistochemistry

Unstained 4 µm lung and TBLN sections were deparaffinized with Propar20, rehydrated with ethanol, and underwent antigen retrieval in citrate buffer before immunofluorescent staining. Slides were UV-bleached to quench autofluorescence, blocked with 5% normal donkey serum and 0.3% Triton X in PBS, then incubated with primary antibodies (B220, CD4, Iba1). After incubation with secondary antibodies, nuclei were counterstained with Hoechst (1:1000). Tissue sections were prepared and stained by the Microscopy and Histology Core Facilities (LJI, La Jolla, CA)

### Evaluation of arthritis and joint damage

Paw and ankle thickness was measured using a caliper (General Ultra Tech TM, No. 1433). Swelling was defined as the increase in thickness from baseline values recorded at the start of the experiment, as described by Mountz et al. [27]. In addition, mice were visually assessed for joint redness, gait and activity changes. For histological assessment of synovitis and bone erosion, paraffin sections (4 μm) of ankles were prepared by the Microscopy and Histology Core Facilities (LJI, La Jolla, CA). Inflammatory arthritis was scored under blinded conditions for synovitis and bone erosion in ankle joints, on a scale of 0-3, as described by Hayer et al.[30].

### Evaluation of development of glomerulonephritis

Periodic Acid–Schiff (PAS) stained paraffin sections (4 μm) of kidney were prepared by the Microscopy and Histology Core Facilities (LJI, La Jolla, CA). Glomerulonephritis was scored on a 0-4 scale under blinded conditions, as described by Koh et al. [31].

### Total immunoglobulin G quantification

Total serum IgG was measured by ELISA as specified by the manufacturer (Immunology Consultants Laboratory, Portland, OR).

### Antinuclear antibodies (ANA) by Indirect Immunofluorescence

Serum antinuclear antibodies (ANA) were detected as previously described [32] using a 1:100 dilution of serum on HEp2 ANA slides (Innova Diagnostics, San Diego, CA). Digital images of ANA patterns were captured using a Leica DFC 365 FX camera and analyzed using Leica Application Suite AF software (Leica Microsystems, Buffalo Grove, IL).

### Autoantibody quantification by ELISA

Quantitative measurement of ENA6-antibodies (anti-Sm, -RNP, -SS-A (60kDa and 52kDa), Jo-1, -SS-B and -Scl-70) and anti-CCP3 was performed using ELISA (QUANTA Lite ELISA Inova Diagnostics, San Diego, CA) modified to detect murine samples using a goat-anti mouse IgG HRP-conjugated antibody (Invitrogen #34137). The units for each sample were calculated as described previously [17]. IgM Rheumatoid Factor (RF) was measured using a sandwich ELISA, as described previously [17].

### High-Throughput Protein Microarray autoantibody Profiling

High-throughput profiling of IgG autoantibodies in serum against various autoantigens (AAgs) was performed by the Microarray and Immune Phenotyping Core Facility, UTSW (Dallas, TX). Briefly, serum samples were pre-treated with DNAse I to remove free-DNA, diluted at 1:50, and hybridized to antigen-coated protein arrays (see Supplementary Table 1). Antibody-binding was detected with Cy3-conjugated anti-mouse IgG (1:2000, Jackson ImmunoResearch Laboratories, Westgrove, PA). Fluorescent images were captured (Genepix 4200A scanner, Molecular Devices, CA) and transformed to signal intensity values using GenePix 7.0 software. Images were background-subtracted and normalized to internal controls. The processed signal intensity value for each autoantibody was reported as antibody score (Ab-score), which is expressed based on the normalized signal intensity (NSI) and signal-to-noise ratio (SNR) using the formula: Ab−score=log2(NSI∗SNR+1). Ab-score values were represented in a heatmap, organized by unsupervised hierarchical clustering generated using the Heatmap.2 package in R. Significant differences between PBS- and c-silica-exposed mice were assessed using t-tests, accounting for multiple testing. Robust autoantibody responses were defined as Ab scores exceeding the PBS average plus one standard deviation (SD).

### Serum inflammatory markers

Serum were tested for inflammatory cytokines and chemokines Rantes/Ccl5, M-CSF, G-CSF, IFN-γ, MIG/Cxcl9, IL-2, IL-5, IL-6, IL-17A, MIP-1β/Ccl4 by MULTIPLEX Map assay according to the manufacturer’s instructions (Millipore Sigma, Burlington, MA).

### Statistical analysis

Data is expressed as mean ± SEM unless otherwise stated. Statistical analysis was performed using GraphPad Software V6 (San Diego, CA). Mann-Whitney U tests were used to compare PBS and c-silica exposed mice. p<0.05 was considered significant. Levels of significance were further indicated as follows: *p<0.05, **p<0.01, ***p<0.001 and ****p<0.0001.

## Results

### C-silica exposure preferentially accelerates development of arthritis over glomerulonephritis in BXD2 mice, but does not induce arthritis or glomerulonephritis in DR4-Tg mice

Arthritis in BXD2 mice was significantly exacerbated by c-silica exposure, as evidenced by pronounced ankle joint swelling (Figure 1A), synovial inflammation and hyperplasia (∼3-fold vs PBS mice) (Figure 1B), and pannus formation with bone and cartilage erosions (∼7-fold vs PBS mice) (Figure 1C) at 20 weeks post-exposure compared to PBS controls. In addition, bone marrow hyperplasia of normal bone marrow cells, consistent with lymphogenesis, was observed (Figure 1F). Minimal arthritis was observed at earlier time points, with no bone erosions detected (data not shown). Two-way ANOVA showed no significant sex or interaction effects, indicating a similar response to silica in both sexes (Figure 1D). In contrast, DR4-Tg mice did not develop inflammatory arthritis at 20 or 40 weeks post-exposure, as evidenced by very limited synovitis (Figure 1E) and absence of bone erosions (data not shown). Representative histological images of ankle (Figure 1F) and knee and front paw (Supplementary Figure 1) illustrate these differences.

**Figure 1:**
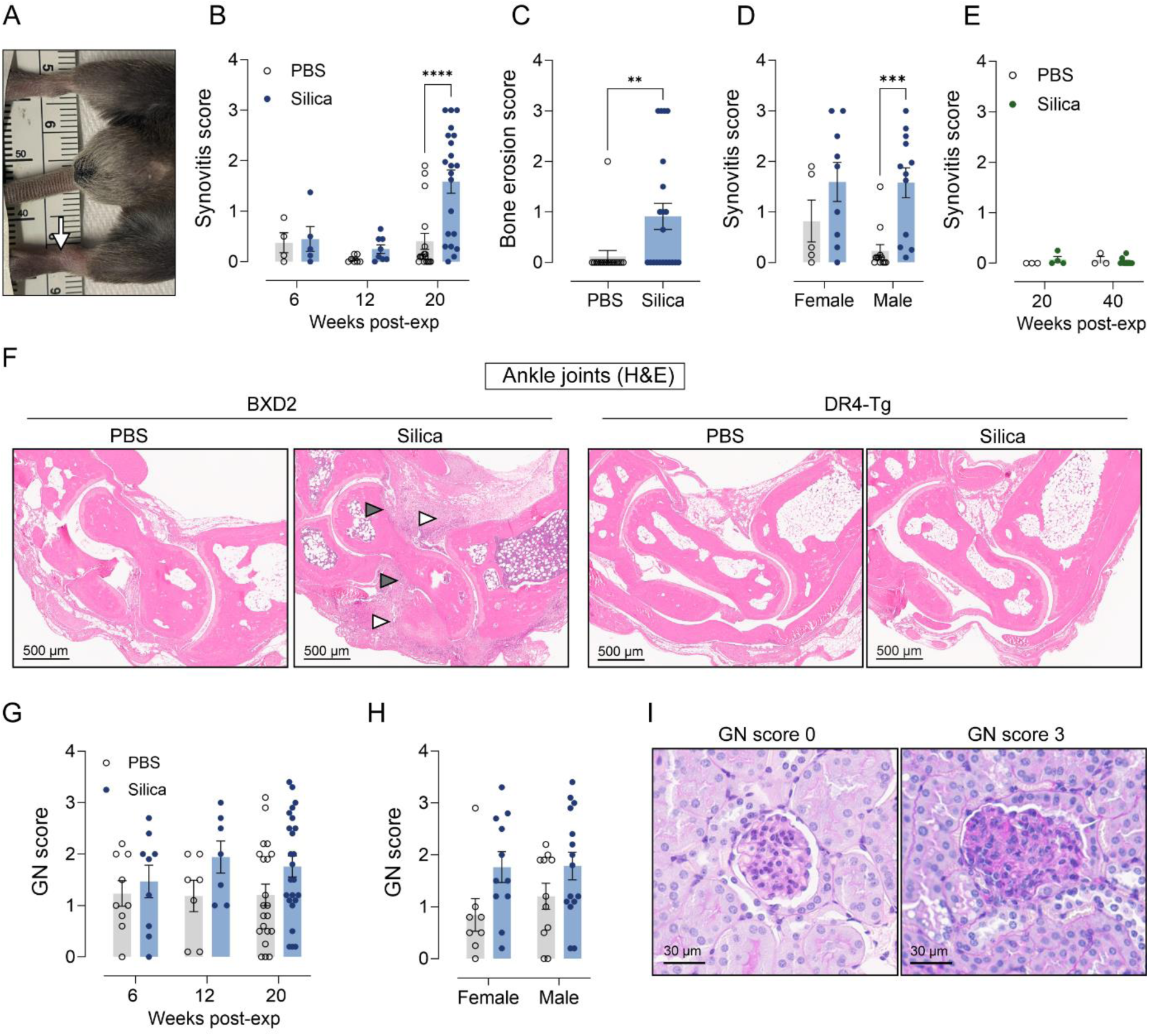
Inflammatory arthritis and glomerulonephritis assessment in BXD2 and DR-Tg mice. (A) Ankles of c-silica-exposed BXD2 mouse with visual swelling of left ankle (arrow). (B) Ankle synovitis scores in c-silica- and PBS-exposed BXD2 mice, 6 (n = 4-5/group), 12 (n = 7-8/group) and 20 weeks post-exposure (n = 17-21/group). (C) Ankle bone erosion scores in c-silica- and PBS-exposed BXD2 mice, 20 weeks post-exposure (n = 17-21 mice/group). (D) Comparison of synovitis scores between female and male c-silica- and PBS-exposed BXD2, 20 weeks post-exposure (n = 5-12 mice/group). (E) Ankle synovitis scores in c-silica- and PBS-exposed DR4-Tg mice, 20 (n = 3-4/group) and 40 weeks post-exposure (n = 3-9/group). (F) Representative histology images of PBS- and c-silica-exposed BXD2 mice, 20 weeks post-exposure, and DR4-Tg mice, 40 weeks post-exposure. Arrowheads indicate synovial inflammation (synovitis) (white arrowheads) and bone erosion/pannus infiltration (grey arrowheads). (G) GN scores from c-silica- and PBS-exposed BXD2 mice at 6, 12 and 20 weeks post-exposure (n = 7-24/group). (H) Comparison of GN scores between female and male c-silica- and PBS-exposed BXD2 mice at 20 weeks post-exposure (n = 8-15/group). (I) Representative kidney tissue images (PAS) of BXD2 mouse that received a GN score of 0 and a BXD2 mouse that received a GN score of 3. Comparisons were performed with Mann-Whitney U tests. *P<0.05, **p<0.01, ***p<0.001, ****p<0.0001.

Kidney histology assessments at 6, 12, and 20 weeks post-exposure revealed no significant differences in glomerulonephritis (GN) severity between c-silica- and PBS-exposed BXD2, though average scores were slightly higher with c-silica. Mild to severe GN was observed in both treatment groups (Figure 1G, representative histology images in Figure 1I). No sex effect was observed (Figure 1H). No GN was observed in DR4-Tg mice (data not shown).

Thus, c-silica exposure in BXD2 predominantly exacerbated inflammatory arthritis and not kidney disease. Moreover, the DR4-Tg mouse model did not develop arthritis or GN and was used in further analyses as a control.

### C-silica exposure induces differential humoral autoimmune features in BXD2 and DR4-Tg mice

To evaluate whether the inflammatory arthritis was accompanied by an increase in generalized autoimmunity in BXD2 but not DR4-Tg mice, spleen weight, lupus and RA autoantibodies, and autoimmunity-associated cytokines and chemokines were assessed.

The systemic autoimmune response induced by c-silica, which was significantly more severe in BXD2 mice than in DR4-Tg mice, was further validated by the observation of c-silica-induced splenomegaly. This was evident in BXD2 mice at 12 and 20 weeks post-exposure (Figure 2A) and in DR4-Tg mice at 40 weeks (Figure 2B). Moreover, total serum IgG levels were significantly higher in c-silica-exposed BXD2 mice vs PBS mice by 6 weeks (Figure 2C), whereas DR4-Tg mice displayed delayed and lower IgG increases compared to BXD2 mice independent of treatment (Figure 2D). Serum ANA levels in BXD2 mice remained unaffected by c-silica exposure, as they were present prior to exposure and increased similarly across groups (Figure 2E). DR4-Tg mice developed lower and delayed ANA responses (Figure 2F).

**Figure 2:**
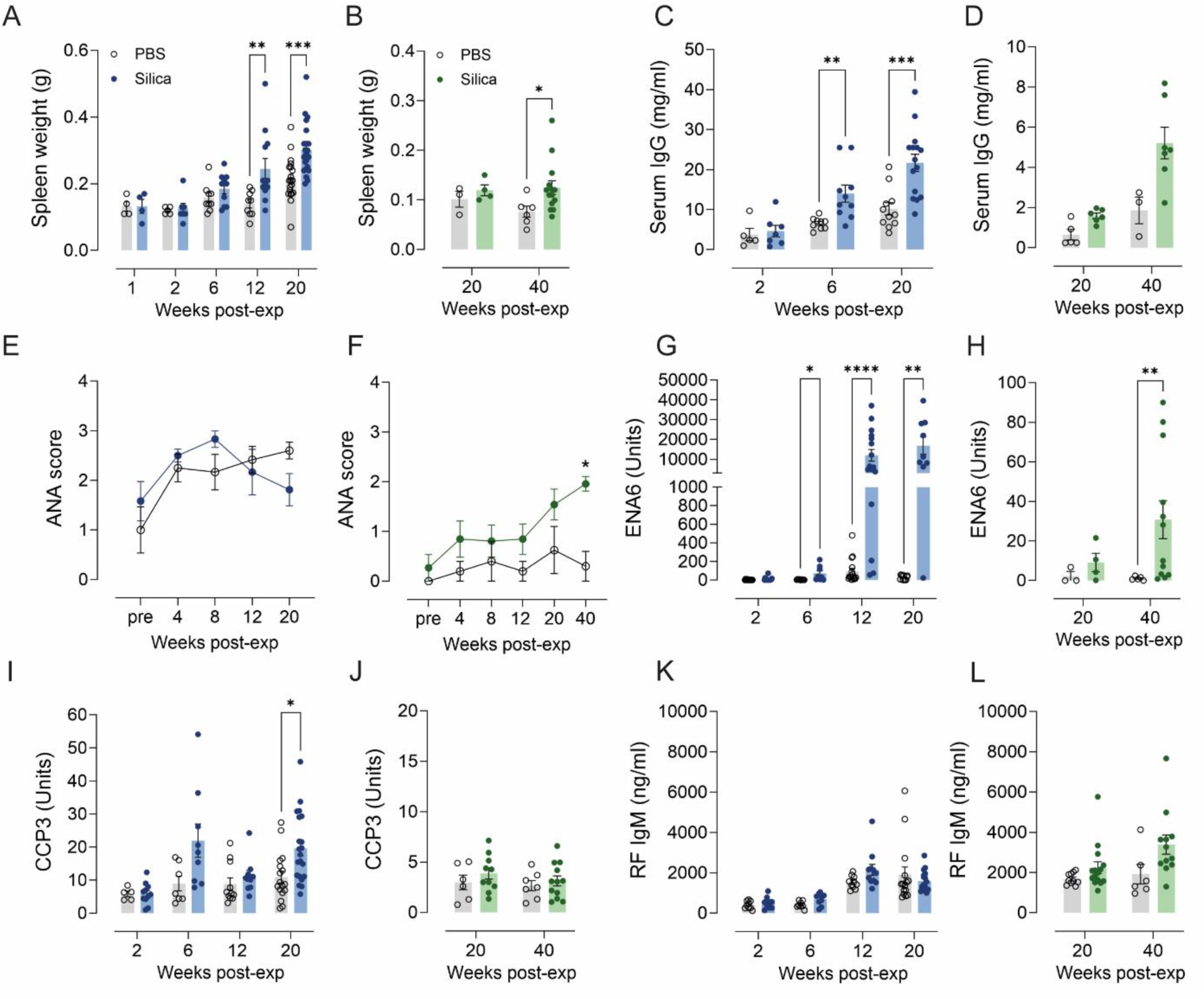
Spleen weight and, total serum IgG and autoantibodies upon c-silica exposure in BXD2 and DR4-Tg mice. Blue graphs represent BXD2 data, green graphs represent DR4-Tg data. Spleen weights from c-silica- and PBS-exposed BXD2 (n = 4-20/group) (A) and DR4-Tg (n = 3-14/group) (B) mice. Total serum IgG (mg/ml) in c-silica- and PBS-exposed BXD2 (n= 5-15/group) (C) and DR4-Tg mice (n = 3-7/group) (D) mice. Serum ANA scores of serum at different timepoints post-exposure in c-silica- and PBS-exposed BXD2 (n = 4-5/group) (E) and DR4-Tg mice (n = 3-8/group) (F). ENA6 levels (units) in serum of c-silica- and PBS-exposed BXD2 (n = 8-17/group) (G) and DR4-Tg mice (n = 3-12/group) (H). CCP3 levels (units) in serum of c-silica- and PBS-exposed BXD2 (n = 5-21/group) (I) and DR4-Tg mice (n = 6-12/group) (J). RF IgM levels in serum (ng/ml) of c-silica- and PBS-exposed BXD2 (n = 10-17/group) (K) and DR4-Tg mice (n = 6-15/group) (L). Groups were compared using Mann-Whitney U tests. *P<0.05, **p<0.01, ***p<0.001, ****p<0.0001.

Autoantigen array analysis performed to assess the diversity of IgG autoantibodies, revealed broad autoantibody repertoires in c-silica exposed BXD2 and DR4-Tg mice. Principal component analyses (PCA) showed almost full separation by treatment in the BXD2 mice, but not the DR4-Tg mice (Figure 3A). The PCA showed a clear separation by strain. C-silica appeared to not only induce the classical autoantibodies, but almost all autoantibody specificities. Co-clustering grouped autoantigens into three main clusters (Figure 3B): the first cluster comprised antigens predominantly targeted in DR4-Tg mice compared to BXD2 mice; the second cluster consisted of autoantigens primarily targeted in c-silica-exposed BXD2 mice; and the third, largest cluster included autoantigens against which BXD2 mice had autoantibodies in both PBS and c-silica-exposed conditions, with these responses further amplified by c-silica exposure. When looking at the strongest and most consistent responses, c-silica exposure of BXD2 mice led to preferential targeting of Sm/RNP antigens, vimentin and anti-ribosomal (P0 and P2) proteins (Figure 3C), while DR4-Tg responses were more diverse, including responses directed toward histones (Figure 3D). These differences were not evident in PBS-treated mice, although DR4-Tg mice generally had lower responses than BXD2 mice (Figure 3E).

**Figure 3:**
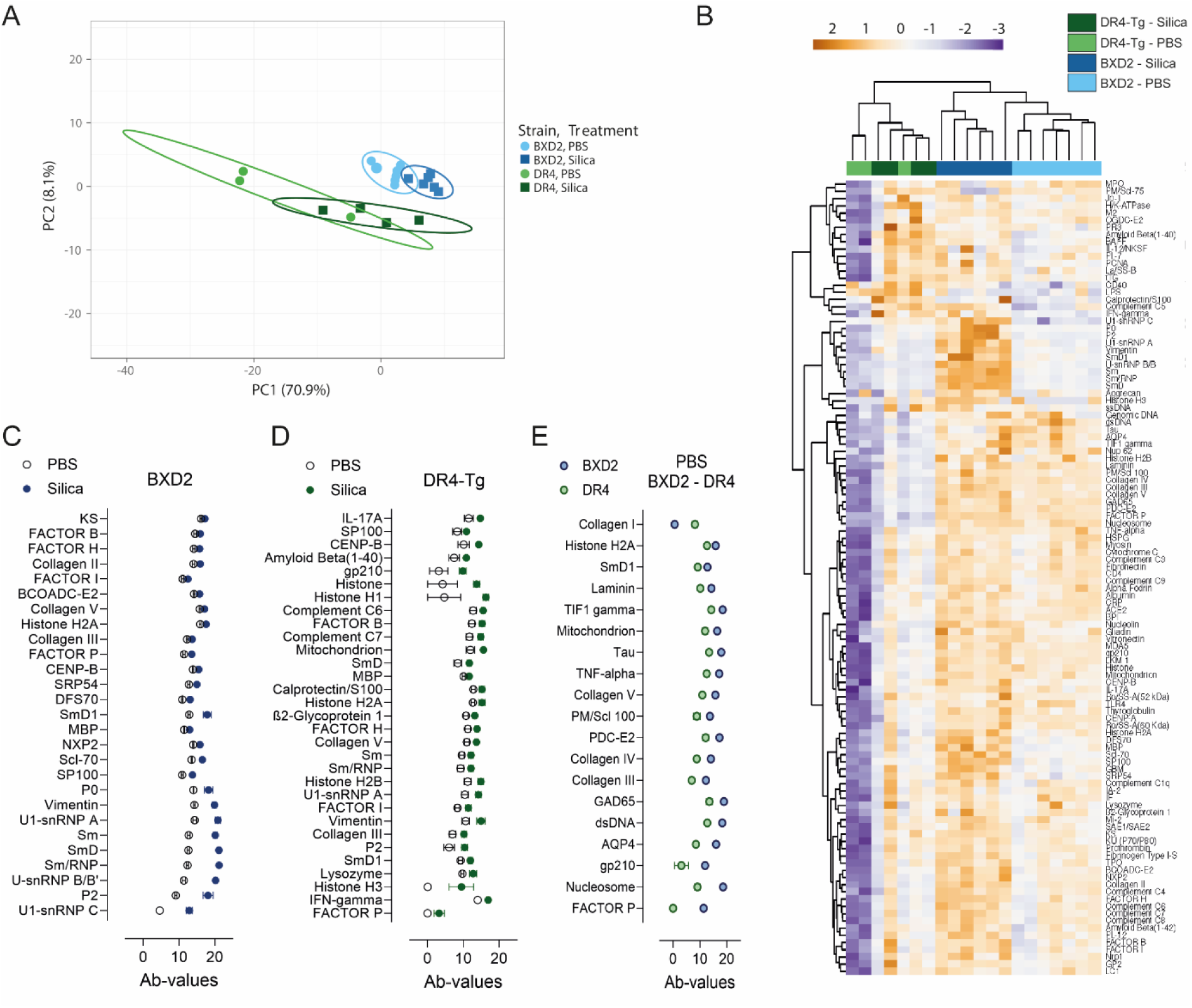
Autoantigen array analyses of serum autoantibodies upon c-silica exposure in BXD2 and DR4-Tg mice. (A) PCA plot and (B) heatmap with unsupervised clustering depict Ab-score values for IgG autoantibody responses against an array of autoantigens in serum samples of c-silica and PBS exposed BXD2 (n = 7-8 mice/group) and DR4-Tg mice (n = 3-4 mice/group), 20 weeks post-exposure. (C) Ab-values of most consistent responses in the BXD2 mice. (D) Ab-values of most consistent responses in the DR4-Tg mice. (E) Ab-values of the responses significantly different between PBS mice of the two strains.

To confirm the strong c-silica-induced response against Sm/RNP antigens, which are part of the extractable nuclear antigens (ENA), we tested the ENA response at different timepoints. Additionally, as responses against citrullinated proteins and rheumatoid factor (IgM) are a hallmark of RA, and as those were not part of the antigen array, we tested these responses at different timepoints. Antibodies against extractable nuclear antigens (ENA6), containing a number of snRNP antigens, were dramatically higher in the c-silica-exposed BXD2 group at 20 weeks (Figure 2G). C-silica-exposed DR4-Tg mice achieved elevated ENA6 responses at 40 weeks though levels remained significantly lower than in BXD2 mice (Figure 2H). Anti-cyclic citrullinated protein (CCP3), antibodies against citrullinated proteins, were significantly elevated at 20 weeks in c-silica-exposed BXD2 mice (Figure 2I), but unexpectedly not increased in DR4-Tg mice after c-silica-exposure (Figure 2J). IgM RF levels were unaffected in BXD2 (Figure 2K) and DR4-Tg mice (Figure 2L).

As autoimmunity is also accompanied by the development of chemokines and cytokines, and as they are known to be elevated by c-silica exposure in bronchioalveolar lavage fluid (BALF) and serum [17, 33], inflammatory markers relevant for autoimmune disease were measured in serum of BXD2 and DR4-Tg mice. Direct comparisons between treatments of the different inflammatory markers showed significant elevations of IL-6, MIG (CXCL9), and MIP-1β (CCL4) in BXD2 mice exposed to c-silica compared to PBS controls, 20 weeks post-exposure (data not shown). In contrast, no significant increase in serum cytokine levels was observed in c-silica-exposed DR4-Tg mice, which also had overall lower levels, even in PBS controls, compared to BXD2. Despite these differences, heatmapping with co-clustering showed only limited clustering based on treatment (c-silica/PBS) and strain (Supplementary Figure 2).

Thus, humoral immune responses in c-silica-exposed BXD2 mice occurred earlier and were greater in magnitude than those of DR4-Tg mice, suggesting significant associations between the humoral autoimmunity and arthritis. Notably, autoantibody responses appeared to be more strongly driven by c-silica exposure than cytokine elevations in serum, highlighting distinct mechanisms of systemic autoimmunity in these strains.

### Pulmonary c-silica exposure triggers development of T- and B-cell containing tertiary lymphoid structures in BXD2, but not DR4-Tg mice

Transoral exposure of c-silica results in pulmonary inflammation characterized by alveolar, peribronchial, and perivascular inflammatory cell infiltrates, and TBLN enlargement [17]. These early inflammatory events can be strain-specific [34], but it is not known if they correlate with systemic responses such as arthritis.

In BXD2 mice, significant peribronchial and perivascular infiltrates were observed at 6 weeks post-exposure, while alveolitis was already present at 1 week (Figure 4A, 4C). DR4-Tg mice exhibited similar alveolitis scores at 20 weeks (Figure 4D) but significantly lower perivasculitis and peribronchitis scores at 20 and 40 weeks (p = 0.004, p = 0.0079; Figure 4B). This resulted in overall higher total lung scores in c-silica-exposed BXD2 mice compared to DR4-Tg mice (Figure 4E, 4F). TBLN enlargement in c-silica-exposed BXD2 mice occurred by 2 weeks post-exposure, progressing over time (Figure 4G), whereas TBLN weights of c-silica-exposed DR4-Tg mice showed less TBLN enlargement at 20 weeks (p = 0.0138; Figure 4H) with delayed increases by 40 weeks. These findings suggest that genetic predisposition in BXD2 drives heightened pulmonary inflammation after c-silica exposure.

**Figure 4:**
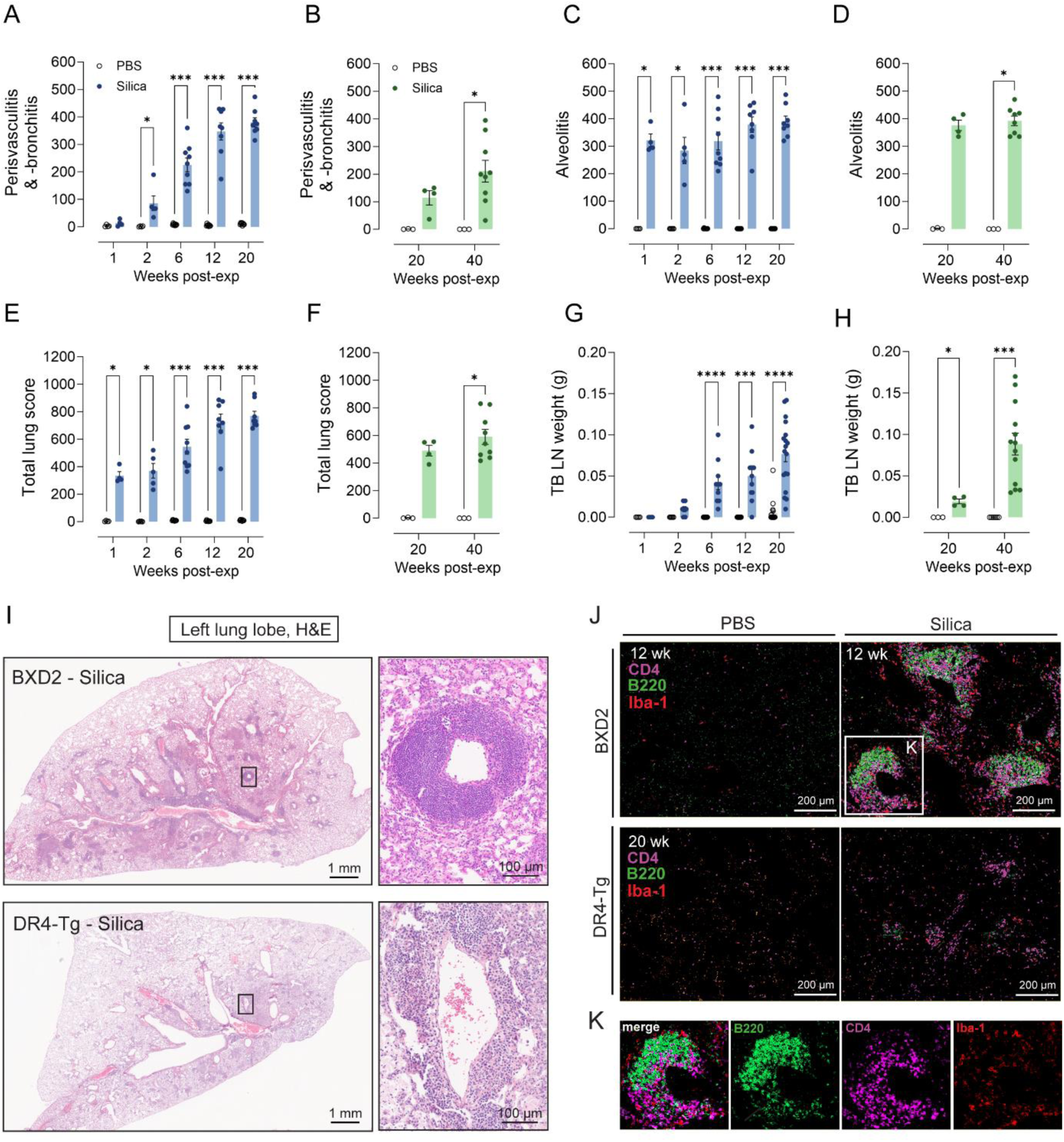
C-silica-induced lung inflammation and iBALT in BXD2 and DR4-Tg mice. Blue graphs represent BXD2 data, green graphs represent DR4-Tg data. Lungs of c-silica- and PBS-exposed BXD2 (n = 4-9/group) and DR4-Tg (n = 3-8/group) mice were scored for alveolitis and peribronchitis and perivasculitis, and scores were added to give a total lung score. Alveolitis score in BXD2 (A) and DR4-Tg mice (B). Peribronchitis and perivasculitis score in BXD2 (C) and DR4-Tg (D) mice. Total lung scores in BXD2 (E) and DR4-Tg (F) mice. Tracheobronchial lymph nodes (TB LN) were collected and weighed in BXD2 mice (n = 4-19 mice/group) (G) and DR4-Tg mice (n = 3-13 mice/group) (H). (I) Representative histology images of H&E-stained lung tissue of c-silica-exposed BXD2 mice, 12 weeks post-exposure, and of c-silica-exposed DR4-Tg mice, 20 weeks post-exposure, showing organized cell accumulations around vasculature and bronchi, present in a higher extent in BXD2 mice compared to DR4-Tg mice. (J) Immunofluorescent staining for CD4, B220 and Iba-1 in lung tissue of c-silica and PBS-exposed BXD2 and DR4-Tg mice shows the formation of iBALT/lymphoid accumulations in BXD2 mice, which are lacking in DR4-Tg mice. (K) Individual markers in lymphoid accumulations from region selected in J. Statistical comparisons were performed with Mann-Whitney U tests. *P<0.05, **p<0.01, ***p<0.001, ****p<0.0001.

Chronic inflammation in non-lymphoid tissues can induce tertiary lymphoid structures (TLS) consisting of organized accumulations of lymphoid cells [35] that are often associated with more severe disease [36]. TLS formation in the lungs, especially near bronchi, termed inducible bronchus-associated lymphoid tissue (iBALT), has been found in RA patients with pulmonary complications, suggesting a relationship with RA pathogenesis [37]. To identify TLS, lung sections were assessed by both H&E and cell-specific immunofluorescent staining. BXD2 mice developed organized lymphoid accumulations resembling iBALT lining vessels and bronchi at 20 weeks post c-silica-exposure (Figure 4I). These TLS contained CD4+ T cells at 2 weeks, progressing to segregated CD4+ and B220+ zones by 6 weeks (Figure 4J, Supplementary Figure 3). By contrast, DR4-Tg mice showed minimal to no TLS formation even at 20 weeks (Figure 4J). In BXD2 mice, iBALT formation coincided with increased perivasculitis and peribronchitis, TBLN enlargement, and autoantibody production (Figures 2, 4A, 4G), linking TLS to severe systemic autoimmunity, including inflammatory arthritis.

### Pulmonary c-silica exposure leads to autoantibodies in BAL fluid of BXD2 mice

Pulmonary inflammation and iBALT or TLS are associated with autoantibodies in sputum and BALF in RA [37, 38] and with autoantibodies in c-silica-induced autoimmunity in mice [19, 39]. To study the relationship of c-silica-induced pulmonary inflammation and iBALT with autoantibody production, anti-ENA6, anti-CCP3, and RF levels in the BALF of c-silica- and PBS-exposed BXD2 and DR4-Tg mice were measured (Figure 5). In BXD2 mice, c-silica exposure significantly increased BALF levels of autoantibodies against ENA6, CCP3, and RF (IgM) compared to PBS controls (Figures 5A, 5C, 5E). BALF autoantibody levels increased over time paralleling the development of perivasculitis and peribronchitis, TBLN, iBALT, and serum autoantibodies (Figures 2, 4). By contrast, DR4-Tg mice exhibited negligible BALF autoantibody levels and no significant changes after c-silica exposure (Figures 5B, 5D, 5F), reflecting reduced pulmonary inflammation and absence of iBALT (Figure 3). These findings highlight the critical role of genetic susceptibility in driving the link between pulmonary inflammation, TLS formation, and systemic autoimmunity.

**Figure 5:**
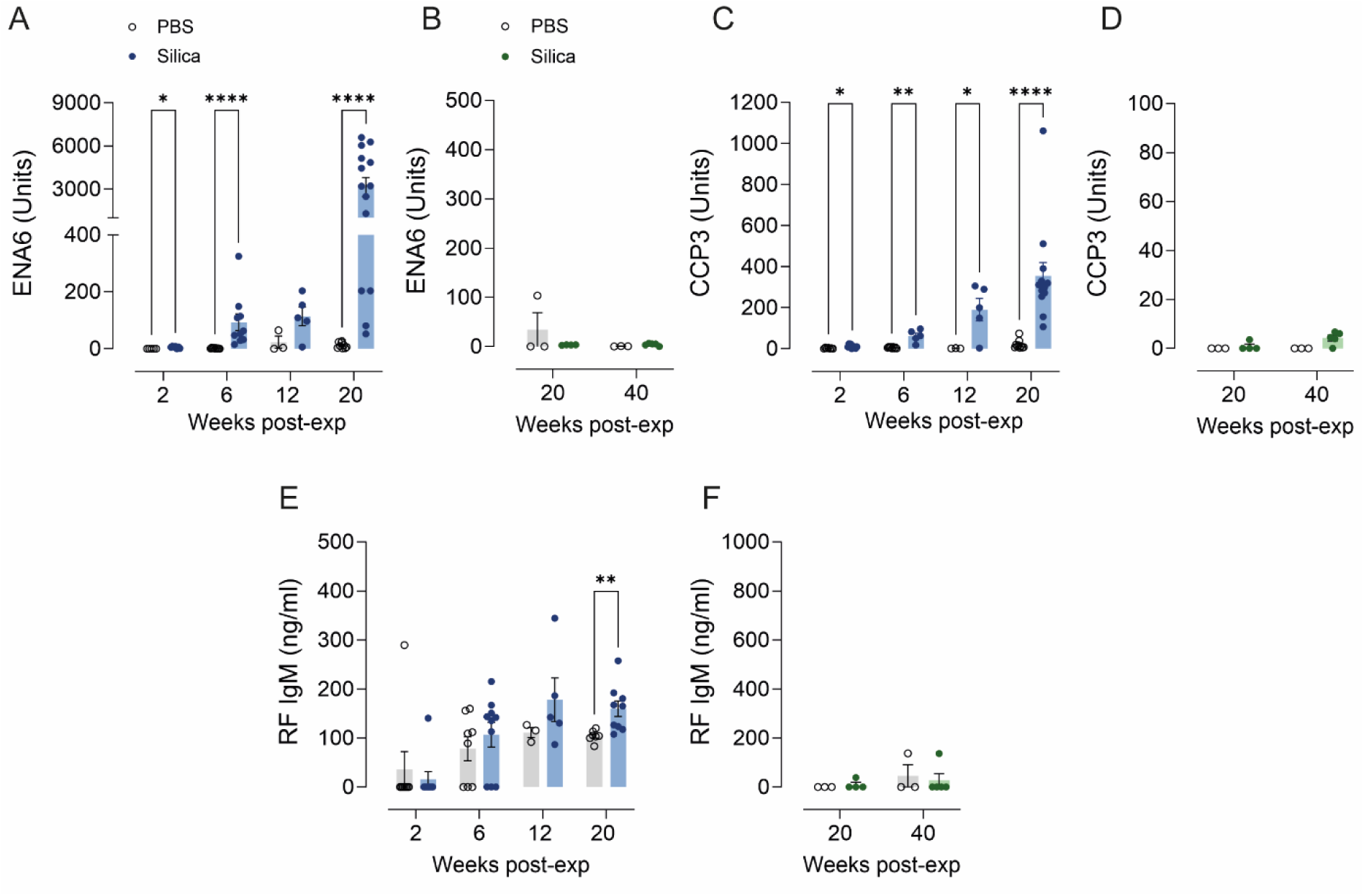
Autoantibodies in BALF of c-silica- and PBS-exposed BXD2 and DR4-Tg mice. Blue graphs represent BXD2 data, green graphs represent DR4-Tg data. Autoantibodies (anti-ENA6, anti-CCP3 and RF IgM) were measured in BALF of c-silica- and PBS-exposed BXD2 mice (n = 3-14 mice/group) and DR4-Tg mice (n = 3-5 mice/group). ENA6 in BXD2 (A) and DR4-Tg mice (B), CCP3 in BXD2 (C) and DR4-Tg (D) mice, and RF IgM in BXD2 (E) and DR4-Tg (F) mice. Statistical comparisons were performed with Mann-Whitney U tests. *P<0.05, **p<0.01, ***p<0.001, ****p<0.0001.

## Discussion

Here we describe a novel experimental model demonstrating that c-silica exposure at an extra-articular site, specifically the lungs, can precipitate inflammatory arthritis in genetically susceptible hosts. Leveraging established models of genetic predisposition to arthritis and RA-associated autoantibodies, we provide new insights into the relationship between inhalant exposure and RA pathogenesis, substantiating the well-documented epidemiological association between occupational c-silica exposure and RA [21]. Our findings further reveal that c-silica induced pronounced pulmonary inflammation in BXD2 mice, characterized by lymphoid cell accumulations at vascular and bronchial sites resembling iBALT. This was accompanied by a robust autoantibody response, notably against snRNP antigens, preceding the exacerbation of synovitis and erosive arthritis. In contrast, c-silica exposure in HLA-DR4 transgenic mice, a strain with a significant genetic risk for RA, failed to induce arthritis and exhibited less pulmonary inflammation, iBALT formation, and autoantibody responses. These findings provide compelling evidence linking pulmonary inflammation, iBALT, and autoantibodies triggered by mucosal environmental exposure to the severity of inflammatory arthritis.

The recombinant inbred (RI) BXD2 line derived from C57BL/6J and DBA/2J strains, exhibits an autoimmune phenotype encompassing erosive arthritis, glomerulonephritis (GN), and autoantibodies including RF and anti-DNA [27, 40]. While BXD2 mice develop features resembling RA, their phenotype extends beyond RA to resemble “rhupus,” a hybrid of RA and lupus [41]. Despite this, the model faithfully recapitulates key RA characteristics, including inflammatory erosive arthritis and RA-like autoantibodies. Mountz et al [27] reported arthritis in BXD2 mice afflicting 50% by 8 months and 90% after 12 months. In our study, evidence of arthritis with visible signs of joint swelling, changes in gait, or erythema were rarely observed in PBS control mice examined around 6 – 7.5 months of age. Nevertheless, histological features of arthritis arising in c-silica exposure mice of the same ages were similar to those described [27], including synovial cell hyperplasia, pannus formation, and severe erosion of cartilage and bone. A greater female prevalence of arthritis was also reported for BXD2 mice similar to idiopathic RA [42], but a sex dichotomy was not observed following c-silica exposure. This is consistent with the absence of sex bias in humans [13] for autoimmune diseases, including RA, associated with c-silica exposure.

The absence of arthritis in c-silica-exposed DR4-Tg mice suggests that apart from MHC Class II genes, other genetic requirements for inflammatory arthritis were missing. This is supported by generally low levels of autoantibodies in serum and BALF. The lack of a c-silica-induced ACPA (anti-CCP) response was unexpected given that other strains that do not expresses the DR4-Tg are capable of making ACPA – we have observed ACPA responses in c-silica exposed Diversity Outbred (DO) mice (Mayeux et al, unpublished data) – and that c-silica exposure increases the ACPA response in mice with CIA [43]. Moreover, DR4-Tg mice can produce ACPA responses when immunized with either citrullinated antigen [23] or type II collagen [44], with both experimental approaches leading to arthritis. HLA-DR4 is thought to promote ACPA by enhancing T cell recognition of citrullinated proteins providing T help for B cells targeting citrullinated proteins [45]. However, it appears that c-silica exposure did not result in an ACPA response in DR4-Tg mice, nor in the development of arthritis.

Although arthritis was significantly accelerated and exacerbated by c-silica in BXD2 mice, this was not the case with GN. Proteinuria in naïve BXD2 mice has been shown to increase at approximately 6 months of age [27]. We found GN, based on kidney histology, to be present as early as 3-4 months of age in some mice but the severity changed little over time or with c-silica exposure, arguing that c-silica does not significantly exacerbate GN in BXD2 mice. However, it is possible that GN occurred before or soon after c-silica exposure, making it difficult to assess acceleration or exacerbation by c-silica exposure. C-silica does accelerate proteinuria and increases the severity of nephritis in lupus-prone NZBWF1 mice in a dose dependent manner [19]. This disparity in strain sensitivity to GN may be reflected by the fact that arthritis and proteinuria can be segregated in F2 mice generated by mating of the (BXD2 X DBA/2)F1 or (BXD2 X B6)F1 mice [27], suggesting a complex genetic control of the arthritis and nephritis phenotypes in BXD2 mice. Moreover, it is possible that c-silica induced pulmonary inflammation promotes immunological and inflammatory factors that favor arthritis pathogenesis more than GN in the BXD2 mice. Only a few of the cytokines typically associated with SLE were elevated by c-silica exposure, possibly explaining the lack of c-silica-induced exacerbation of GN.

Although our studies do not provide a causal mechanism explaining the importance of pulmonary inflammation to the development of synovitis and bone erosion, we can conclude that localized inflammation in the lungs precedes development of autoantibodies and that the increasing severity of inflammation and autoimmunity is important in the subsequent acceleration of inflammatory arthritis. A significant observation is the development of TLS with features of iBALT in the lungs following c-silica exposure. Such well-defined structures have also been found in lupus-prone NZBWF1 mice exposed to c-silica [19] but, to our knowledge, have not been formally described in humans with c-silica exposure. TLS, also called tertiary lymphoid organs or ectopic lymphoid structures, can be the result of a response against antigenic stimuli such as infection or environmental insult [46]. iBALT has been described in patients with RA [37] and similar structures have been found in RA joints [47] and the kidney in SLE [48], suggestive of a significant role in disease pathology. It is unclear whether these structures amplify autoantibody responses or initiate autoantibody production, but it is clear that they are linked to the presence of autoantibodies, including ACPA and RF [37, 46, 49]. Pulmonary inflammation in RA is argued to lead to protein citrullination and local production of ACPA [49, 50]. Because of the close temporal relationship between pulmonary inflammation, TBLN enlargement, iBALT, and serum autoantibody levels following c-silica exposure, our studies do not establish whether the initiation of the autoantibody response begins in iBALT or the TBLN. However, in naïve mice, the small size of the TBLN makes it difficult to locate but it rapidly increases in size following c-silica exposure and develops c-silica containing nodules [17]. This suggests the trafficking of antigenic material from the lung to the TBLN which may lead to T and B cell activation and expansion and migration to inflammatory sites in the lung. Analysis of the antigenic specificity of B cells in both locations over time is needed to establish when and where c-silica-induced autoreactive B cell responses begin.

An important biomarker of RA is the presence of disease-related autoantibodies with ACPA and RF being the most significant [51], both of which are found in RA linked to c-silica exposure [9, 25, 52]. Although the presence of such antibodies can precede clinical diagnosis [53], their role in pathogenesis is unclear [50, 54]. However, antibodies have been found to induce arthritis in experimental models, particularly anti-collagen II and anti-G6PI antibodies [55]. BXD2 mice had elevations of both ACPA (anti-CCP3) and RF in BALF and ACPA in serum while DR4-Tg mice had elevation of RF in serum only. Autoantibody responses were low in BALF, likely reflecting the lower levels of immunoglobulin in BALF compared to serum. However, unlike anti-ENA6, anti-CCP3 levels were also low in serum arguing that anti-CCP3 may originate in the lung. The most prominent autoantibody response was directed against ENA6, a mix of Sm, RNP, SS-A (60kDa and 52kDa), SS-B, Scl-70, and -Jo-1. Previous studies [16, 17] as well as the antigen array analysis used here, identify Sm and RNP as the major targets of the c-silica-induced autoantibody response. Using a similar antigen array, Rajasinghe et al [56] found a diverse spectrum of responses against nuclear, non-nuclear, and tissue specific antigens in BALF and serum of lupus-prone NZBWF1 mice exposed to c-silica. While our studies also found a diverse array of antigenic targets in BXD2 mice, the greatest discrimination between PBS controls and c-silica exposed mice was against polypeptides that are part of Sm and RNP snRNPs. Similar autoantibody responses have been found in those occupationally exposure to c-silica dust including responses against centromeric proteins, Scl70 (topoisomerase I), DNA, as well as RNA binding proteins Ro(60)/SSA and La/SSB and the snRNP component U1-RNP [57]. Thus, a common feature of the c-silica-induced autoantibody response in humans and experimental models is against nucleic acid, especially RNA, binding proteins. Although the reason for this is unknown, it may be related to the ability of RNA binding proteins from cell lysates to coat silica nanoparticles [58] which could potentially provide a source of antigenic material following phagocytosis of c-silica particles. To our knowledge, little has been done to define the autoantibody profile in RA patients occupationally exposed to silica.

Our findings lend robust experimental support to the mucosal origins hypothesis of RA, which posits that inflammatory events at mucosal surfaces, such as the lungs, precede systemic autoimmunity and joint inflammation [49]. Unlike prior rodent models that relied on direct joint manipulation or systemic immunization [43, 59], this study uniquely employs mucosal exposure to c-silica as the sole arthritis trigger, recapitulating key features of RA, including genetic susceptibility, mucosal inflammation, autoantibody production, and erosive arthritis. This study establishes a novel experimental framework connecting occupationally relevant c-silica exposure to RA pathogenesis. By demonstrating the interplay between pulmonary inflammation, iBALT formation, and systemic autoimmunity in the progression of inflammatory arthritis, our findings bridge epidemiological data and mechanistic insights. Future research should explore the cellular and molecular pathways linking mucosal inflammation to distant joint pathology, further illuminating the enigmatic relationship between environmental exposures and autoimmune diseases.

## Acknowledgements

We thank the Microscopy and Histology Core Facilities of the La Jolla Institute for Immunology, La Jolla California, and the Microarray and Immune Phenotyping Core Facility at The University of Texas Southwestern Medical Center, Dallas, TX for their excellent services.

## Author contributions

Conceptualization: KMP, DHK, JM, LMFJ; Experiments: LMFJ, JM, CDO; Histology scoring: DHK; Writing-Original Draft Preparation: LMFJ; Writing-Review and Editing: KMP, DHK, CDO, MG, PHMH, JM; Visualization: LMFJ; Supervision: KMP, DHK, MG, SR, PHMH, JM; Funding Acquisition: KMP.

## References

1. England B.R., Baker J.F., Sayles H., et al., Body Mass Index, Weight Loss, and Cause-Specific Mortality in Rheumatoid Arthritis. Arthritis Care & Research, 2018. 70(1): p. 11–18.

2. England B.R., Sayles H., Michaud K., et al., Cause-Specific Mortality in Male US Veterans With Rheumatoid Arthritis. Arthritis Care & Research, 2016. 68(1): p. 36–45.

3. Kelly C.A., Saravanan V., Nisar M., et al., Rheumatoid arthritis-related interstitial lung disease: associations, prognostic factors and physiological and radiological characteristics-a large multicentre UK study. Rheumatology, 2014. 53(9): p. 1676–1682.

4. Silman A.J. and Pearson J.E., Epidemiology and genetics of rheumatoid arthritis. Arthritis Research & Therapy, 2002. 4: p. S265–S272.

5. Gregersen P.K., Goyert S.M., Song Q.L., et al., Microheterogeneity of HLA-DR4 haplotypes: DNA sequence analysis of LD“KT2” and LD“TAS” haplotypes. Hum Immunol, 1987. 19(4): p. 287–92.

6. Gregersen P.K., Silver J., and Winchester R.J., The Shared Epitope Hypothesis - an Approach to Understanding the Molecular-Genetics of Susceptibility to Rheumatoid-Arthritis. Arthritis and Rheumatism, 1987. 30(11): p. 1205–1213.

7. Deighton C.M., Walker D.J., Griffiths I.D., et al., The Contribution of Hla to Rheumatoid-Arthritis. Clinical Genetics, 1989. 36(3): p. 178–182.

8. Padyukov L., Silva C., Stolt P., et al., A gene-environment interaction between smoking and shared epitope genes in HLA-DR provides a high risk of seropositive rheumatoid arthritis. Arthritis Rheum, 2004. 50(10): p. 3085–92.

9. Stolt P., Yahya A., Bengtsson C., et al., Silica exposure among male current smokers is associated with a high risk of developing ACPA-positive rheumatoid arthritis. Annals of the Rheumatic Diseases, 2010. 69(6): p. 1072–1076.

10. Mehri F., Jenabi E., Bashirian S., et al., The association Between Occupational Exposure to silica and Risk of Developing Rheumatoid Arthritis: A Meta-Analysis. Saf Health Work, 2020. 11(2): p. 136–142.

11. Pollard K.M., Silica, Silicosis, and Autoimmunity. Frontiers in Immunology, 2016. 7.

12. Leung C.C., Yu I.T.S., and Chen W.H., Silicosis. Lancet, 2012. 379(9830): p. 2008–2018.

13. Boudigaard S.H., Schlünssen V., Vestergaard J.M., et al., Occupational exposure to respirable crystalline silica and risk of autoimmune rheumatic diseases: a nationwide cohort study. International Journal of Epidemiology, 2021. 50(4): p. 1213–1226.

14. Parks C.G., Miller F.W., Pollard K.M., et al., Expert panel workshop consensus statement on the role of the environment in the development of autoimmune disease. Int J Mol Sci, 2014. 15(8): p. 14269–97.

15. Parks C.G., Conrad K., and Cooper G.S., Occupational exposure to crystalline silica and autoimmune disease. Environ Health Perspect, 1999. 107 **Suppl 5**(Suppl 5): p. 793–802.

16. Gonzalez-Quintial R., Mayeux J.M., Kono D.H., et al., Silica exposure and chronic virus infection synergistically promote lupus-like systemic autoimmunity in mice with low genetic predisposition. Clin Immunol, 2019. 205: p. 75–82.

17. Mayeux J.M., Escalante G.M., Christy J.M., et al., Silicosis and Silica-Induced Autoimmunity in the Diversity Outbred Mouse. Front Immunol, 2018. 9: p. 874.

18. Brown J.M., Archer A.J., Pfau J.C., et al., Silica accelerated systemic autoimmune disease in lupus-prone New Zealand mixed mice. Clin Exp Immunol, 2003. 131(3): p. 415–21.

19. Bates M.A., Brandenberger C., Langohr I., et al., Silica Triggers Inflammation and Ectopic Lymphoid Neogenesis in the Lungs in Parallel with Accelerated Onset of Systemic Autoimmunity and Glomerulonephritis in the Lupus-Prone NZBWF1 Mouse. PLoS One, 2015. 10(5): p. e0125481.

20. Janssen L.M.F., Ghosh M., Lemaire F., et al., Exposure to silicates and systemic autoimmune-related outcomes in rodents: a systematic review. Particle and Fibre Toxicology, 2022. 19(1).

21. Miller F.W., Alfredsson L., Costenbader K.H., et al., Epidemiology of environmental exposures and human autoimmune diseases: findings from a National Institute of Environmental Health Sciences Expert Panel Workshop. J Autoimmun, 2012. 39(4): p. 259–71.

22. Zhao T., Xie Z., Xi Y., et al., How to Model Rheumatoid Arthritis in Animals: From Rodents to Non-Human Primates. Front Immunol, 2022. 13: p. 887460.

23. Hill J.A., Bell D.A., Brintnell W., et al., Arthritis induced by posttranslationally modified (citrullinated) fibrinogen in DR4-IE transgenic mice. J Exp Med, 2008. 205(4): p. 967–79.

24. Klareskog L., Stolt P., Lundberg K., et al., A new model for an etiology of rheumatoid arthritis: smoking may trigger HLA-DR (shared epitope)-restricted immune reactions to autoantigens modified by citrullination. Arthritis Rheum, 2006. 54(1): p. 38–46.

25. Yahya A., Bengtsson C., Larsson P., et al., Silica exposure is associated with an increased risk of developing ACPA-positive rheumatoid arthritis in an Asian population: evidence from the Malaysian MyEIRA case-control study. Mod Rheumatol, 2013.

26. Mohamed B.M., Verma N.K., Davies A.M., et al., Citrullination of proteins: a common post-translational modification pathway induced by different nanoparticles in vitro and in vivo. Nanomedicine (Lond), 2012. 7(8): p. 1181–95.

27. Mountz J.D., Yang P., Wu Q., et al., Genetic segregation of spontaneous erosive arthritis and generalized autoimmune disease in the BXD2 recombinant inbred strain of mice. Scand J Immunol, 2005. 61(2): p. 128–38.

28. Hamilton R.F., Jr., Thakur S.A., Mayfair J.K., et al., MARCO mediates silica uptake and toxicity in alveolar macrophages from C57BL/6 mice. J Biol Chem, 2006. 281(45): p. 34218–26.

29. Mayeux J.M., Kono D.H., and Pollard K.M., Development of experimental silicosis in inbred and outbred mice depends on instillation volume. Sci Rep, 2019. 9(1): p. 14190.

30. Hayer S., Vervoordeldonk M.J., Denis M.C., et al., ’SMASH’ recommendations for standardised microscopic arthritis scoring of histological sections from inflammatory arthritis animal models. Ann Rheum Dis, 2021. 80(6): p. 714–726.

31. Koh Y.T., Scatizzi J.C., Gahan J.D., et al., Role of nucleic acid-sensing TLRs in diverse autoantibody specificities and anti-nuclear antibody-producing B cells. J Immunol, 2013. 190(10): p. 4982–90.

32. Toomey C.B., Cauvi D.M., Hamel J.C., et al., Cathepsin B regulates the appearance and severity of mercury-induced inflammation and autoimmunity. Toxicol Sci, 2014. 142(2): p. 339–49.

33. Rajasinghe L.D., Bates M.A., Benninghoff A.D., et al., Silica Induction of Diverse Inflammatory Proteome in Lungs of Lupus-Prone Mice Quelled by Dietary Docosahexaenoic Acid Supplementation. Front Immunol, 2021. 12: p. 781446.

34. Davis G.S., Leslie K.O., and Hemenway D.R., Silicosis in mice: effects of dose, time, and genetic strain. J Environ Pathol Toxicol Oncol, 1998. 17(2): p. 81–97.

35. Dong Y., Wang T., and Wu H., Tertiary lymphoid structures in autoimmune diseases. Front Immunol, 2023. 14: p. 1322035.

36. Sato Y., Silina K., van den Broek M., et al., The roles of tertiary lymphoid structures in chronic diseases. Nat Rev Nephrol, 2023. 19(8): p. 525–537.

37. Rangel-Moreno J., Hartson L., Navarro C., et al., Inducible bronchus-associated lymphoid tissue (iBALT) in patients with pulmonary complications of rheumatoid arthritis. J Clin Invest, 2006. 116(12): p. 3183–94.

38. Demoruelle M.K., Wilson T.M., and Deane K.D., Lung inflammation in the pathogenesis of rheumatoid arthritis. Immunol Rev, 2020. 294(1): p. 124–132.

39. Fee L., Kumar A., Tighe R.M., et al., Autoreactive B cells recruited to lungs by silica exposure contribute to local autoantibody production in autoimmune-prone BXSB and B cell receptor transgenic mice. Front Immunol, 2022. 13: p. 933360.

40. Hamilton J.A., Li J., Wu Q., et al., General Approach for Tetramer-Based Identification of Autoantigen-Reactive B Cells: Characterization of La- and snRNP-Reactive B Cells in Autoimmune BXD2 Mice. J Immunol, 2015. 194(10): p. 5022–34.

41. Panush R.S., Edwards N.L., Longley S., et al., ’Rhupus’ syndrome. Arch Intern Med, 1988. 148(7): p. 1633–6.

42. Pollard K.M., Gender differences in autoimmunity associated with exposure to environmental factors. J Autoimmun, 2012. 38(2-3): p. J177–86.

43. Engelmann R. and Muller-Hilke B., Experimental silicosis does not aggravate collagen-induced arthritis in mice. J Negat Results Biomed, 2017. 16(1): p. 5.

44. Taneja V., Behrens M., Mangalam A., et al., New humanized HLA-DR4-transgenic mice that mimic the sex bias of rheumatoid arthritis. Arthritis Rheum, 2007. 56(1): p. 69–78.

45. Hill J.A., Southwood S., Sette A., et al., Cutting edge: The conversion of arginine to citrulline allows for a high-affinity peptide interaction with the rheumatoid arthritis-associated HLA-DRB1*0401 MHC class II molecule. Journal of Immunology, 2003. 171(2): p. 538–541.

46. Marin N.D., Dunlap M.D., Kaushal D., et al., Friend or Foe: The Protective and Pathological Roles of Inducible Bronchus-Associated Lymphoid Tissue in Pulmonary Diseases. J Immunol, 2019. 202(9): p. 2519–2526.

47. Rivellese F., Pontarini E., and Pitzalis C., Tertiary Lymphoid Organs in Rheumatoid Arthritis. Curr Top Microbiol Immunol, 2020. 426: p. 119–141.

48. Jamaly S., Rakaee M., Abdi R., et al., Interplay of immune and kidney resident cells in the formation of tertiary lymphoid structures in lupus nephritis. Autoimmun Rev, 2021. 20(12): p. 102980.

49. Holers V.M., Demoruelle M.K., Kuhn K.A., et al., Rheumatoid arthritis and the mucosal origins hypothesis: protection turns to destruction. Nat Rev Rheumatol, 2018. 14(9): p. 542–557.

50. Gravallese E.M. and Firestein G.S., Rheumatoid Arthritis - Common Origins, Divergent Mechanisms. New England Journal of Medicine, 2023. 388(6): p. 529–542.

51. Sokolove J., Wagner C.A., Lahey L.J., et al., Increased inflammation and disease activity among current cigarette smokers with rheumatoid arthritis: a cross-sectional analysis of US veterans. Rheumatology, 2016. 55(11): p. 1969–1977.

52. Stolt P., Källberg H., Lundberg I., et al., Silica exposure is associated with increased risk of developing rheumatoid arthritis:: results from the Swedish EIRA study. Annals of the Rheumatic Diseases, 2005. 64(4): p. 582–586.

53. Greenblatt H.K., Kim H.A., Bettner L.F., et al., Preclinical rheumatoid arthritis and rheumatoid arthritis prevention. Curr Opin Rheumatol, 2020. 32(3): p. 289–296.

54. Sokolova M.V., Schett G., and Steffen U., Autoantibodies in Rheumatoid Arthritis: Historical Background and Novel Findings. Clin Rev Allergy Immunol, 2022. 63(2): p. 138–151.

55. Caplazi P., Baca M., Barck K., et al., Mouse Models of Rheumatoid Arthritis. Vet Pathol, 2015. 52(5): p. 819–26.

56. Rajasinghe L.D., Li Q.Z., Zhu C., et al., Omega-3 fatty acid intake suppresses induction of diverse autoantibody repertoire by crystalline silica in lupus-prone mice. Autoimmunity, 2020. 53(7): p. 415–433.

57. Conrad K. and Mehlhorn J., Diagnostic and prognostic relevance of autoantibodies in uranium miners. Int Arch Allergy Immunol, 2000. 123(1): p. 77–91.

58. Klein G., Mathe C., Biola-Clier M., et al., RNA-binding proteins are a major target of silica nanoparticles in cell extracts. Nanotoxicology, 2016. 10(10): p. 1555–1564.

59. Nowak B., Majka G., Srottek M., et al., The Effect of Inhaled Air Particulate Matter SRM 1648a on the Development of Mild Collagen-Induced Arthritis in DBA/J Mice. Arch Immunol Ther Exp (Warsz), 2022. 70(1): p. 17.

