## Supplementary Materials for "Silica-mediated exacerbation of inflammatory arthritis: A novel murine model"

**Supplementary File 1:** Animal number information.

**Supplementary Figure 1:** Knee joint and front paw histology images (H&E) of c-silica- and PBS-exposed BXD2 mice.

**Supplementary Figure 2:** Heatmap of cytokine and chemokine values (pg/ml) in serum of c-silica- and PBS-exposed BXD2 and DR4-Tg mice.

**Supplementary Figure 3:** Immunofluorescent staining (CD4, B220 and Iba1) of lung tissue of c-silica-exposed BXD2 mice at 1, 2, 6 and 12 weeks post-exposure.

### Supplementary Figures + legends

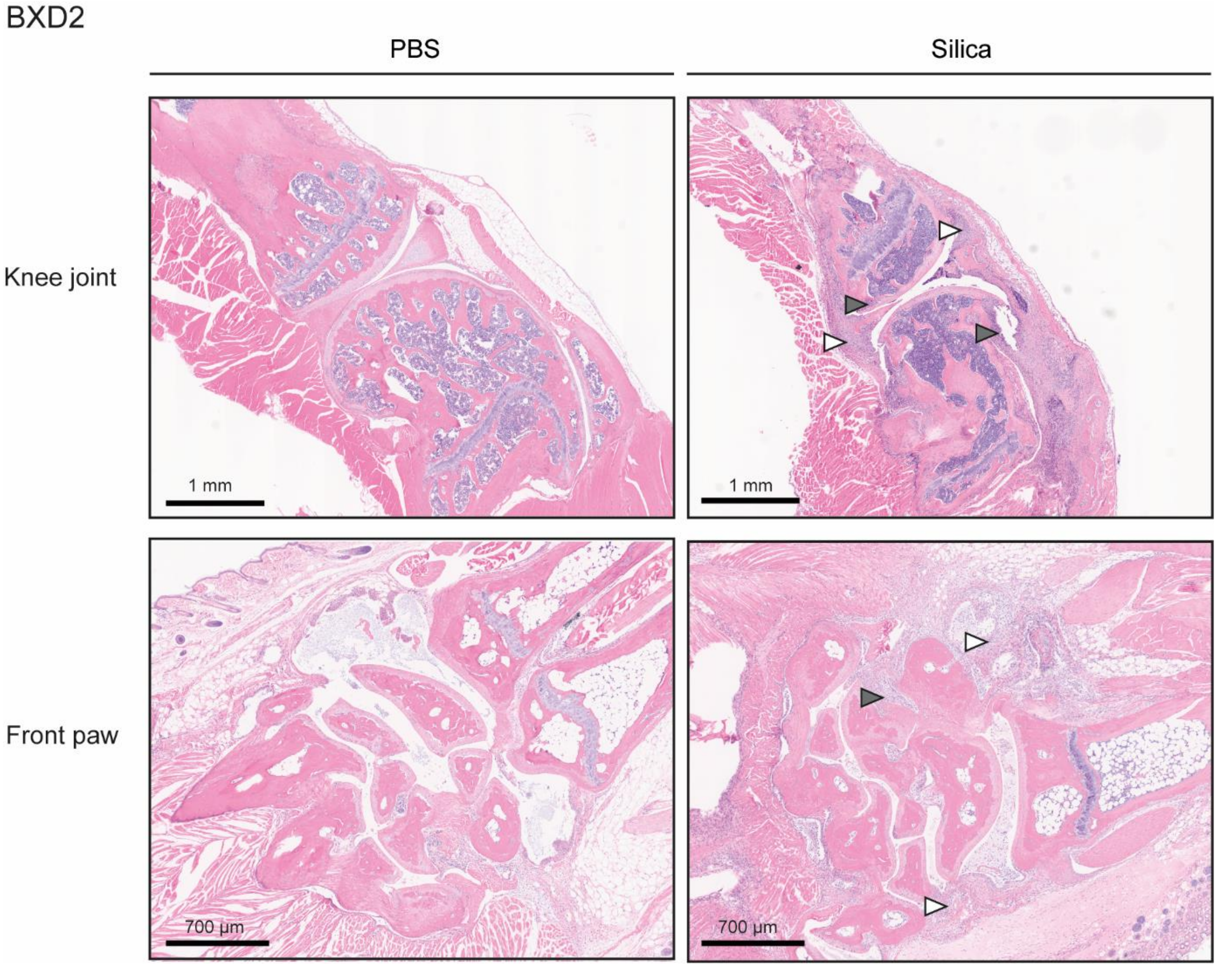


**Supplementary Figure 1:** Knee joint and front paw histology images (H&E) of c-silica- and PBS-exposed BXD2 mice. White arrows indicate synovitis, grey arrows indicate bone erosion.

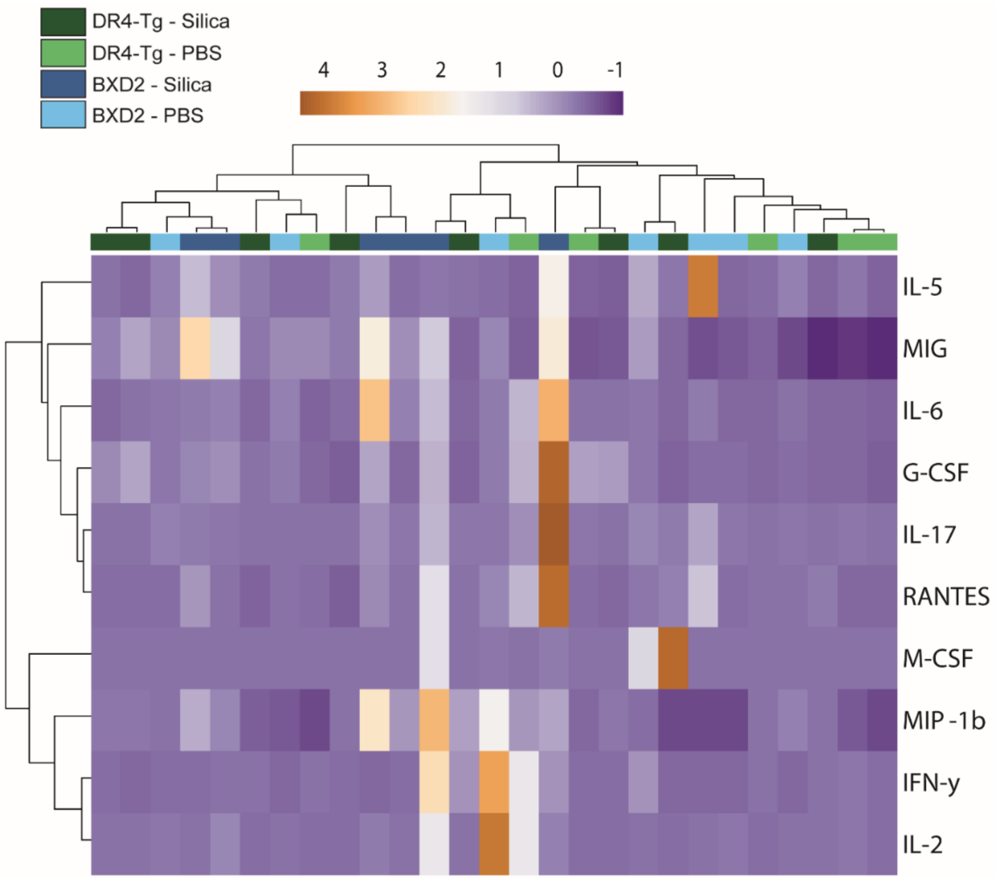


**Supplementary Figure 2:** Heatmap of cytokine and chemokine values (pg/ml) in serum of c-silica- and PBS-exposed BXD2 and DR4-Tg mice (20 weeks post-exposure). Heatmapping with co-clustering was performed on Z-scores of raw values (pg/ml).

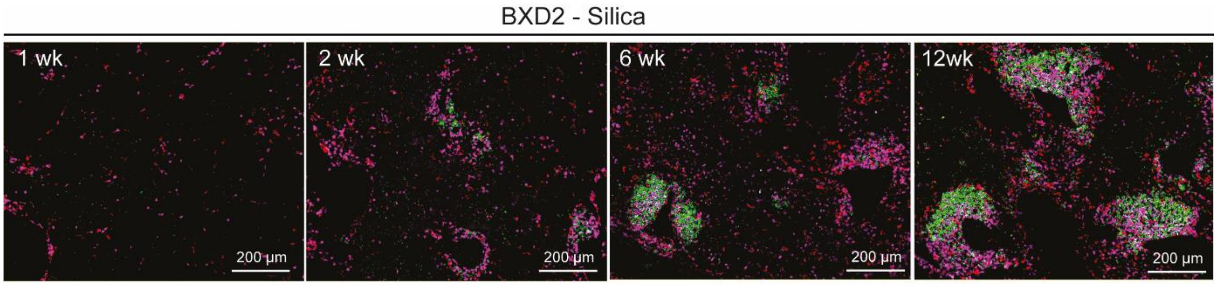


**Supplementary Figure 3:** Immunofluorescent staining (CD4, B220 and Iba1) of lung tissue of c-silica-exposed BXD2 mice at 1, 2, 6 and 12 weeks post-exposure.

### Supplementary File 1

#### Mouse harvest numbers.

The table below shows the number of mice harvested per cohort. The sample size for each cohort is based on historical mean values and standard deviations observed in previous published studies indicating that 4 mice provides 80% power to detect 30% change at a significance level of 0.05 and standard deviation of 30%, and is sufficient for detecting clinically relevant changes in the immunopathologic parameters that were measured including autoantibodies, histology, immunohistochemistry, and inflammatory markers in silica-exposed mice. This size acted as a default if no other relevant information was available. The sample size for each cohort was further determined based on several factors, including mouse availability, attrition due to mortality during the experiment, and variability in specific outcomes observed during preliminary studies. Larger cohort sizes were used when we observed higher variability in specific outcomes, to ensure robust statistical analyses and reliable interpretation of the data.

| | BXD2 | | DR4 | |
|---|---|---|---|---|
| Weeks post-exposure | PBS | Silica | PBS | Silica |
| 1 | 4 | 4 | / | / |
| 2 | 9 | 10 | / | / |
| 6 | 8 | 10 | / | / |
| 12 | 9 | 12 | / | / |
| 20 | 17 | 24 | 3 | 4 |
| 40 | / | / | 3 | 9 |

#### Sample Inclusion and Exclusion Criteria

For each measurement and outcome, mice were evaluated based on specific inclusion and exclusion criteria to ensure data reliability and consistency. The criteria applied are as follows:

1. **Overall exclusion criteria:**
    - Suspected infections: Mice, regardless of exposure group, were excluded if they exhibited severe lung inflammation that was inconsistent with expected outcomes and suggested a potential respiratory infection.

- Failed silica exposure: Silica-exposed mice were excluded if histological analysis of the lung tissue failed to detect silica particles by polarized microscopy of tissue, indicating unsuccessful exposure.

2. **Endpoint-specific exclusion criteria**

    - Sample quality: Mice were excluded from specific analyses if the collected tissue did not meet quality standards for the intended assay (e.g., compromised histological integrity).
    - Sample prioritization: If available sample was limited, the sample was not included in each endpoint.

Therefore, n-values are included underneath each figure for each measurement.

**Randomization.**

Treatments (silica or PBS) were assigned randomly within cages, such that mice in the same cage could receive different treatments. No formal randomization method was used.

**Blinding for group allocation.**

Blinding for group allocation was ensured at different levels.

Blinded for group allocation:

- Tissue staining preparation for histology and histology scoring.
- Performing autoantibody and cytokine measurements.
- Performing autoantibody array measurements.
- Performing immunofluorescent staining.

Not blinded for group allocation:

- During silica/PBS treatments.
- During harvest of mice.
